# Multiple mechanisms of aminoglycoside ototoxicity are distinguished by subcellular localization of action

**DOI:** 10.1101/2024.05.30.596537

**Authors:** Patricia Wu, Francisco Barros Becker, Roberto Ogelman, Esra D. Camci, Tor H. Linbo, Julian A. Simon, Edwin W Rubel, David W. Raible

**Author notes:** Correspondence: David W. Raible.

## Abstract

Mechanosensory hair cells of the inner ears and lateral line of vertebrates display heightened vulnerability to environmental insult, with damage resulting in hearing and balance disorders. An important example is hair cell loss due to exposure to toxic agents including therapeutic drugs such as the aminoglycoside antibiotics such as neomycin and gentamicin and antineoplastic agents. We describe two distinct cellular pathways for aminoglycoside-induced hair cell death in zebrafish lateral line hair cells. Neomycin exposure results in death from acute exposure with most cells dying within 1 hour of exposure. By contrast, exposure to gentamicin results primarily in delayed hair cell death, taking up to 24 hours for maximal effect. Washout experiments demonstrate that delayed death does not require continuous exposure, demonstrating two mechanisms where downstream responses differ in their timing. Acute damage is associated with mitochondrial calcium fluxes and can be alleviated by the mitochondrially-targeted antioxidant mitoTEMPO, while delayed death is independent of these factors. Conversely delayed death is associated with lysosomal accumulation and is reduced by altering endolysosomal function, while acute death is not sensitive to lysosomal manipulations. These experiments reveal the complexity of responses of hair cells to closely related compounds, suggesting that intervention focusing on early events rather than specific death pathways may be a successful therapeutic strategy.

## Introduction

The inner ear is essential for normal vocal communication, for localizing acoustic information and for balance. The sensitivity of the ear to disease and injury is evident by the large number of mutations that affect its development and maintenance (Friedman et al., 2007; Angeli et al., 2012), and by the well-documented susceptibility of hair cells to damage from important therapeutic agents such as aminoglycoside (AG) antibiotics and antineoplastic agents including cisplatin (Rizzi and Hirose, 2007; Sheth et al., 2017; Jiang et al., 2017; O’Sullivan et al., 2017). AGs are standard-of-care treatment for life-threatening diseases of the lung, including multidrug-resistant tuberculosis (van Altena et al., 2017; Heysel et al., 2018), nontuberculous mycobacterium (Dillard et al., 2021), and *Pseudomonas* infections associated with cystic fibrosis (Scheenstra et al., 2009; Garinis et al., 2017; 2021; Handelsman et al., 2017; Zettner et al., 2018), with associated hearing loss ranging from 11-67% of patients. In some cases, damage to vestibular organs is even more prevalent than hearing loss (Handelsman et al., 2017). Aminoglycoside use has been estimated to contribute to nearly 20 million cases of hearing loss globally (Prasad et al., 2024). Understanding how AG exposure results in hair cell death may provide potential therapeutic avenues to make these important drugs safer.

The zebrafish lateral line system has emerged as a useful model for identifying hearing-related genes and their functions, and for understanding cellular pathways underlying some forms of hair cell death (reviewed in Stawicki et al., 2015; Coffin and Ramcharitar, 2016; Nicolson, 2017; Kindt and Sheets, 2018; Pickett and Raible, 2019). Mechanosensory hair cells are located on the surface of the body in clusters, called neuromasts (NMs), making them amenable to visualization and manipulation. Like hair cells of the inner ear, lateral line hair cells demonstrate hypersusceptiblity to damage by exposure to toxic chemicals, including AGs and cisplatin (Harris et al., 2003; Ton and Parng, 2005; Ou et al., 2007). The zebrafish lateral line system has proven useful in screening for ototoxic drugs (Chiu et al., 2008; Hirose et al., 2011; Neveux et al., 2017; Davis et al., 2020; Coffin et al., 2021) and screening for potentially protective therapeutics (Owens et al., 2008; Vlasits et al., 2012; Kenyon et al., 2017; Chowdhury et al., 2018).

In zebrafish, intercompartmental calcium (Ca^2+^) flows play a central role in lateral line hair cell toxicity after acute exposure to the AG, neomycin. Treatment with neomycin results in endoplasmic reticulum flow of Ca^2+^ via IP_3_ channels to mitochondria, with subsequent loss of mitochondrial membrane potential (Esterberg et al., 2013; 2014). Ultrastructural analyses of zebrafish hair cells revealed rapid loss of mitochondrial integrity after AG exposures that result in hair cell death (Owens et al., 2007). Increases in mitochondrial Ca^2+^ are accompanied by increases in reactive oxygen species (ROS), and modulating mitochondrial ROS production confers some protection against neomycin damage (Esterberg et al., 2016). Together, these studies suggested a model where neomycin disruption of ER-mitochondrial Ca^2+^ flows resulted in mitochondrial collapse, and cell death. However, it is not clear whether this model is generalizable to other ototoxins, including other AG antibiotics, and previous studies demonstrated differences in temporal response of lateral line hair cells to different AGs (Owens et al., 2009; Coffin et al., 2013), suggesting that there may be multiple pathways by which zebrafish hair cells die from AG exposure.

By varying both AG concentration and time of exposure, we now provide evidence for an acute mechanism that happens within minutes of AG exposure and a delayed mechanism that occurs for hours after AG exposure. The two patterns display distinct intracellular Ca^2+^ dynamics and differential accumulation of AGs in lysosomes. We further demonstrate that altering lysosomal distribution protects against delayed but not acute damage. Together, these studies reveal multiple vulnerabilities of hair cells to AG exposure and suggest that development of therapeutic approaches to preserve hearing and balance need to consider common and distinct pathways underlying damage.

## Methods

### Animals

Experiments were conducted on 5-8 dpf larval zebrafish. Larvae were raised in embryo medium (EM; 14.97 mM NaCl, 500 μM KCL, 42 μM Na2HPO4, 150 μM KH2PO4, 1 mM CaCl2 dihydrate, 1 mM MgSO4, 0.714 mM NaHCO3, pH 7.2) at 28.5°C. All wildtype animals were of the AB strain. Zebrafish experiments and husbandry followed standard protocols in accordance with University of Washington Institutional Animal Care and Use Committee guidelines.

The *Tg(myosin6b:RGECO1)^vo10Tg^* line, expressing the red Ca^2+^ indicator RGECO in the cytoplasmic compartment, has been previously described (Maeda et al., 2014) and was provided as gift from Katie Kindt (National Institute of Deafness and Other Communication Disorders, Bethesda, MD, USA). The *Tg(myosin6b:MITO-GCaMP3)^w119Tg^* line, expressing the green Ca^2+^ indicator GCaMP in the mitochondrial compartment, was described previously (Esterberg et al., 2014). The line *Tg(pou4f3:GAP-GFP)^s273tTg^*expresses membrane-tagged Green Fluorescence Protein (GFP) in hair cells (Xiao et al., 2005). The line *Tg(myosin6b:EGFP-rab7a)^w272Tg^*was generated using the Tol2 transposon (Kawakami et al., 2000) and the hair cell-specific *myo6b* promoter (Kindt et al., 2012) to direct expression of eGFP fused 5’ to the zebrafish *rab7a* gene (Clark et al., 2011).

### AG Treatment

To assess the relative toxicity of AGs, 5-8 dpf larvae were treated with neomycin (Sigma-Aldrich, N1142), gentamicin (Sigma-Aldrich, G1397) or G418 (Sigma-Aldrich, A1720). For acute treatment, larvae were exposed to AG in EM for 1h, and then were rinsed 3x in EM, euthanized with 1.3% MESAB (MS-222; ethyl-m-aminobenzoate methanesulfonate; Sigma-Aldrich, A5040) and fixed (see below) for hair cell analysis. For delayed (1+23h) treatment, larvae were treated for 1h in AG, then rinsed 3x in EM followed by incubation in EM for 23h, euthanized and fixed for hair cell analysis. For 24h treatment, larvae were treated with AG for 24h, and then were rinsed 3x in EM, euthanized and fixed for hair cell analysis.

To test whether subcellular localization influenced toxicity, larvae were first treated with drugs that inhibit lysosomal or mitochondria uptake. We used 250µM GPN (glycyl-l-phenylalanine 2-naphthylamide; Cayman, 14634), or 100 nM Bafilomycin A1 (Sigma-Aldrich, B1793) to alter lysosomal function, or with 50µM mitoTEMPO (Sigma-Aldrich, SML0737) to alter mitochondrial oxidation. Drugs were first added for 1 h (GPN and Bafilomycin) or 30 min (mitoTEMPO) before the addition of AG for one hour.

To assess AG accumulation in hair cells we conjugated G418 or neomycin to the fluorophore Texas Red-X-succinimidyl ester (ThermoFisher, T6134) or G418 to BODIPY 650/665-X NHS Ester (Succinimidyl ester; ThermoFisher, D10001) following protocols for gentamicin labeling (Sandoval et al., 1998; Steyger et al., 2003) with previously described modifications (Stawicki et al., 2014). For lysosomal distribution experiments, groups of 3 5dpf larvae were incubated in embryo media (EM) with 250µM GPN for 1 h. Then, G418-Bodipy650 was added to final concentration of 50µM for 1 h. Larvae were then mounted and imaged (described below). For experiments comparing AG distribution in hair cells, groups of 2 5dpf larvae were imaged at a time. Larvae were exposed to 25µM Neomycin-Texas Red or G418-Texas Red for 5 min in EM. Drug was washed out by transferring larvae to a 60×15mm Petri dish (Falcon, 351007) containing EM+0.2% MESAB for 1min, mounted and imaged (described below).

### Hair cell analysis

Larvae were fixed with 4% paraformaldehyde in 0.1 M phosphate-buffered saline (PBS, pH 7.4) for 1 hr at room temperature (RT) in a slow moving rocker. Larvae were then washed 3x with PBS for 15min at RT and blocked in PBS supplemented with 0.1 % Triton-X 100 and 5% normal goat serum (Sigma-Aldrich, 11H280) for 1-2 hrs at RT. Samples were incubated in mouse anti-parvalbumin antibody (1:400; EMD Millipore, MAB1572) in blocking solution (PBS supplemented with 0.1 % Triton-X 100 and 1% normal goat serum) at 4°C overnight. Then samples were rinsed with PBS-T (PBS, 1% Triton-X 100) and subsequently incubated in goat anti-mouse antibody conjugated to Alexa 488 or 568 (1:500; ThermoFisher, A-11001, A11004) in blocking solution at 4°C overnight. Larvae were rinsed with PBS-T 3x for 15min at RT, then with PBS 3x for 15min at RT, and mounted between coverslips with Fluoromount G (SouthernBiotech, 0100-01). Hair cells were counted using a Zeiss Axioplan 2ie epifluorescence microscope with a Plan-NEOFLUAR 40x/0.75 NA objective (Zeiss). Counts were performed on four neuromasts (NMs) per fish (SO1, SO2, O1, and OC1; Raible and Kruse, 2000) and summed to arrive at one value per fish. Hair cell counts were performed for 9 - 13 fish per treatment condition.

### Calcium Imaging

Calcium imaging was performed on live fish as previously described (Esterberg et al., 2014) using an inverted Marianas spinning disk system (Intelligent Imaging Innovations, 3i) with an Evolve 10 MHz EMCCD camera (Photometrics) and a Zeiss C-Apochromat 63x/1.2 NA water objective. Fish were stabilized using a slice anchor harp (Harvard Instruments) so that neuromasts on immobilized animals had access to the surrounding media. Imaging was performed at ambient temperature, typically 25°C. Baseline fluorescence readings were taken before AG exposure in 30 s intervals for 2.5 min. Aminoglycoside was added as a 4X concentrated stock to achieve the final indicated concentration. Fluorescence intensity readings were acquired in 30 s intervals for 60 min. Camera intensification was set to keep exposure times <50 ms for GCaMP, 250 ms for cytoRGECO while keeping pixel intensity <25% of saturation. Camera gain was set at 3X to minimize photobleaching. Z-sections were taken at 1 μm intervals through the depth of the neuromast, typically 24 μm. GCaMP fluorescence was acquired with 488 nm laser and 535/30 emission filter. RGECO was acquired with a 561 nm laser and a 617/73 emission filter.

For analyses, maximum intensity projections were generated and frames were auto-aligned using SlideBook software (Intelligent Imaging Innovations) to account for XY drift, typically <50 pixels. ROIs outlining the cell of interest were drawn by hand, enabling us to correct for individual cell movement when necessary. Cells were categorized as living or dying based on their clearance from the neuromast. Fluorescence intensities were calculated relative to the mean baseline intensity of each individual hair cell before aminoglycoside exposure. For each treatment condition, at least three replications were performed on different days and fluorescence intensities of no more than three cells per neuromast and two neuromasts per animal were used in analyses. Living and dying cells were chosen randomly for analysis at the end of each time lapse.

### Confocal Image acquisition

Live larvae were anesthetized in EM+0.2% MESAB, and then transferred to a glass-bottom chamber (Ibidi, m-Slide 2) and mounted using a small piece of 1mm x 1mm net, and two slice tissue harps (Warner Instruments, SHD-26GH/10 and SHD-27LH/10). Larvae were imaged using a Zeiss LSM 980 Airyscan inverted microscope, using a LD LCI Plan-Apochromat 40x/1.2NA Imm autocorr DIC objective (Zeiss). Correction collar was set to a relative position of 70% (*Tg(myosin6b:EGFP)*) or 75% (*Tg(myosin6b:EGFP-Rab7a)*). 6x digital zoom was used. Z-stack span was defined for each neuromast, keeping an interval of 0.21 μm. Images were obtained using the Airyscan FAST SR-4Y scanning method, with optimal sampling confirmed by Zen Blue 3.7. The laser line was aligned to the Airyscan sensor array before imaging each larvae group.

### Image processing

3D Airyscan processing of all images was performed on Zen Blue 3.7, with a strength value of 4.5 for all channels. For images of Tg(myosin6b:EGFP-Rab7a) larvae, background fluorescence was subtracted on the EGFP-Rab7a channel using the rolling ball macro in FIJI (Schindelin et al., 2012), with a pixel size of 10. For images of Tg(myosin6b:EGFP) larvae, a custom Python script (Hewitt et al. 2024), was used to align the z-stacks with the pystackreg Python port of StackReg (Thevenaz et al., 1998). Using parameters of ‘rigid body’ as method and ‘previous’ as reference settings, alignment matrices were calculated using the cytoplasmic GFP signal, and then applied to all channels. Alignment matrices, and aligned images were then saved as.OME.tiff files. A third of images were randomly selected and visually examined to make sure alignment was done correctly.

### Image segmentation

We used 3D automated image segmentation to measure the accumulation of labeled AGs into vesicles. To segment the neuromast using the EGFP-Rab7a signal, we applied a strong gaussian filter (sigma value of 6), followed by triangle threshold (Zack et al., 1997) from scikit-image (van der Walt et al., 2014). This mask was processed to fill artificial holes and eliminate small artifacts generated during the segmentation. To create the background mask, we used the inverted neuromast mask. To segment neuromasts and hair cells using the myosin6b:EGFP transgene, we implemented a two-step semi-automated process, with a final user correction step. First, a background mask was created by thresholding using a multi-Otsu function from scikit-image (Liao et al., 2001), with a class value of 5. We then replaced pixel intensity in the background mask with an arbitrary value of 1.0e-7. To identify hair cells, we next created a mask by applying a Sauvola local thresholding (J. Sauvola and M. Pietikäinen, 2000), with a window size of 45 and a k value of 0.2. This mask was processed to fill artificial holes and eliminate small artifacts generated during the segmentation. Individual hair cells were then identified using watershed segmentation. Seeds (one per cell) were added manually using a distance transform map as reference, using a Napari points layer (Sofroniew et al., 2022). Watershed segmentation results were visualized in Napari, and errors in the cell masks were corrected by hand using the draw function.

Vesicles were segmented using the Allen Cell and Structure Segmenter (Chen et al., 2018). Files were loaded and visualized using Napari viewer and the range of slices in the stack that contain the neuromast image were selected. Intensity normalization function was applied to both channels. Vesicles were segmented using the 2D slice-by-slice dot wrapper and 2D filament wrapper functions from the Allen Cell Classic Segmenter package (Chen et al., 2018) with the following parameters: for EGFP-Rab7a vesicles (Figure 5), Dot segmenter parameters: scale_spot_1 = 4, cutoff_spot_1 = 0.5, scale_spot_2 = 2, cutoff_spot_2 = 0.08, and Filament segmenter parameter: scale_fil_1 = 2, cutoff_fil_1 = 1, scale_fil_2 = 1, cutoff_fil_2 = 0.9; for G418 vesicles (Figure 6), Dot segmenter parameters: scale_spot_1 = 4, cutoff_spot_1 = 0.04, scale_spot_2 = 1, cutoff_spot_2 = 0.05, and Filament segmenter parameter: scale_fil_1 = 3, cutoff_fil_1 = 0.2, scale_fil_2 = 2, cutoff_fil_2 = 0.09. The two segmenter results were fused together to create a single vesicle mask. This mask was then processed to fill artificial holes and eliminate small segmentation artifacts. To identify individual vesicles we used the label function from scikit-image (Wu et al., 2005), acknowledging that some vesicle masks might end up labeled together. Any segmented vesicles outside the neuromast mask were discarded for further analysis. A cytoplasmic mask was created by subtracting the vesicular mask from the whole neuromast mask. Properties for each segmented mask were calculated using regionprops_table function from scikit-image.

### Statistical analyses

Mann-Whitney tests, T-tests, one way or two-way ANOVA with posthoc testing was performed using GraphPad Prism. For graphical presentation, data were normalized to untreated controls such that 100% represents hair cell survival in control animals. Results were considered significant if p ≤ 0.05 with levels of statistical significance shown in figures as follows: * 0.01 to 0.05, ** 0.001 to 0.01, *** 0.0001 to 0.001, **** < 0.0001.

## Results

### Distinct mechanisms of hair cell death are revealed by varying the exposure times to AGs

To illustrate differences in susceptibility to damage, we compare hair cell death after exposure to two different AGs, neomycin and gentamicin (Figure 1). After 1h exposure (Figure 1A), there was substantial loss of hair cells with 200 µM neomycin treatment, whereas exposure to the same concentration of gentamicin had little effect. After 24h (Figure 1B), there was substantial damage with both AGs. We considered two hypotheses to account for the difference in gentamicin exposure resulting in these different outcomes. One possibility was that continuous exposure to gentamicin over 24h was required for damage similar to that seen with a comparable concentration of neomycin to occur. Alternatively, the cellular responses to damage after exposure to gentamicin might be delayed, requiring further incubation time but not additional exposure. To distinguish between these ideas, we treated hair cells with AGs for 1h, followed by rinsing in fresh medium to remove all AG, and then incubating for an additional 23h (designated 1+23h treatment). We found that 1h exposure to 200 µM gentamicin was sufficient to cause damage comparable to that seen with neomycin if animals were incubated an additional 23h in fresh medium (Figure 1C). These results are consistent with the idea that 1h gentamicin exposure results in cell death processes that are substantially delayed compared to those that result from 1h neomycin treatment.

**Figure 1.**
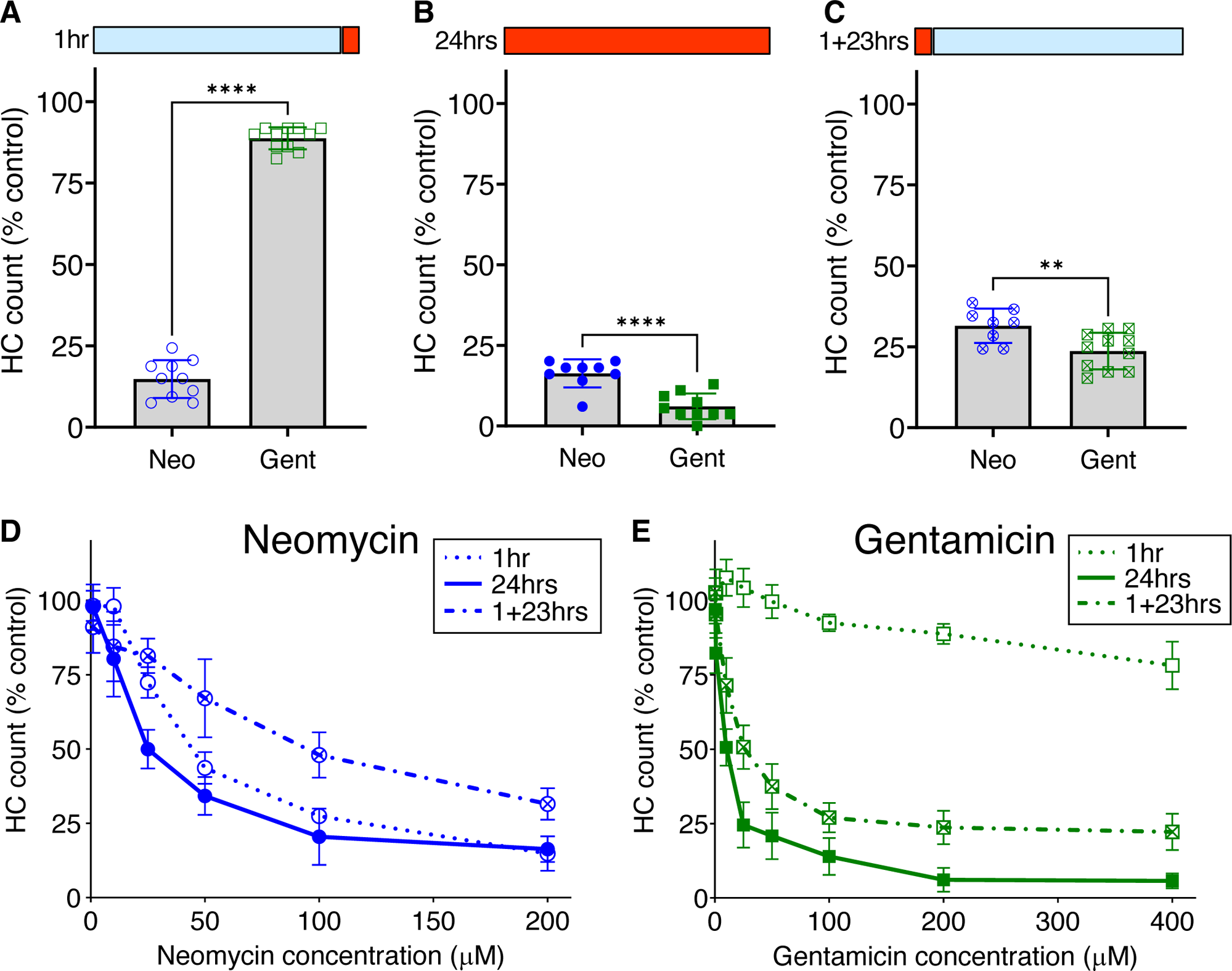
AGs differ in relative hair cell toxicity dependent on time of exposure and length of incubation. (A-C) Fish were treated with 200 µM neomycin (Neo) or gentamicin (Gent) (red bar) for time indicated, or rinsed into fresh medium (blue bar). A) Hair cells were exposed for 1hr acute treatment with AGs. Hair cells are effectively killed by Neo, but are spared by Gent. n=9-11 fish, 4 NMs/fish. Unpaired t test with Welch’s correction, **** P value <0.0001 B) Chronic 24hrs treatment with Neo or Gent results in hair cell loss. n=9-11 fish, 4 NMs/fish. Unpaired t test with Welch’s correction, **** P value <0.0001 C) Delayed death occurs after 1hr treatment with Neo or Gent, rinsing, and then incubation for 23hrs in fresh medium (1+23hrs). n=9-11 fish, 4 NMs/fish. Unpaired t test with Welch’s correction, ** P value = 0.0073 D) Dose-dependent loss of hair cells after treatment with neomycin for 1hr, 24hrs or 1+23hrs. Differences between treatments were highly significant (2-way ANOVA, Tukey’s multiple comparison, p<0.0005). n=9-11 fish, 4 NMs/fish for each condition. E) Dose-dependent loss of hair cells after treatment with gentamicin for 1hr, 24hrs or 1+23hrs. Differences between treatments were highly significant (2-way ANOVA, Tukey’s multiple comparison, p<0.0001). n=9-11 fish, 4 NMs/fish for each condition. Error bars represent Standard Deviation.

To better characterize relative sensitivities, we analyzed dose-response functions of each AG, with concentrations up to 400 µM, using the three experimental paradigms: 1h, 1+ 23h, or 24h exposure. For neomycin (Figure 1D), treatment following all three paradigms resulted in substantial and significant dose-dependent damage (2-way ANOVA: p<0.0001) with significant differences between treatment paradigms (p<0.0001). While there was a statistically distinct increase in sensitivity across the dose-response function from 24h continuous exposure compared to 1h acute exposure (Tukey’s multiple comparison, p<0.0001), both treatment paradigms resulted in substantial dose-dependent damage. There were significantly more hair cells present with the 1+23h paradigm compared to 1h acute treatment (Tukey’s multiple comparison, p=0.0003). This increase is probably due to the rapid initiation of hair cell regeneration after neomycin washout (Harris et al., 2003; Ma et al., 2008). Treatment with gentamicin (Figure 1E) following all three paradigms also resulted in substantial and significant dose-dependent damage (2-way ANOVA: p<0.0001), with significant differences between each of the three treatment paradigms (p<0.0001). While both 24h and 1+23 treatment paradigms resulted in substantial loss of hair cells, there was nevertheless a large and highly significant difference between these treatments (Tukey’s multiple comparison, p<0.0001). By contrast, treatment for 1h caused only a slight but significant loss of hair cells across the dose range (1-way ANOVA; p<0.001). One way to compare relative efficacy of drug treatments is to calculate the concentration where 50% of hair cells are killed, designated HC50 (Chowdhury et al., 2018). For 1h acute treatment with neomycin, the HC50 is 44 µM, compared to 25 µM for 24h treatment and 94 µM for 1+23 treatment. For gentamicin, treatment for 24h was most effective, with an HC50 of 10 µM, and the 1+23h treatment almost as effective with an HC50 of 19 µM. Treatment with gentamicin for 1h never approached 50% kill, with only 20% loss at the highest doses. These results demonstrate that differences in the timing of response of hair cells to different AGs persist across the dose-response function.

We next assessed when hair cells were lost during the 23h incubation period after 1h gentamicin exposure. We exposed larval fish to gentamicin at concentrations ranging from 25 µM to 200 µM for 1h, then rinsed into fresh medium and incubated further for 5, 11, 17 or 23h before quantifying hair cell loss compared to untreated controls. Figure 2 shows a regular decline in the number of hair cells over this time period, demonstrating that cells are gradually lost. We observed significant dose-dependent damage (2-way ANOVA: F=275.9, p<0.0001) with significant differences between treatment paradigms (F=118.2, p<0.0001). The rate of hair cell loss was dependent on the initial concentration of gentamicin present during the 1h exposure period. While the total hair cell loss assessed at the 24h endpoint was similar across initial concentrations, hair cell loss varied greatly at intermediate times in a manner dependent on concentration. Treatment with 25 µM gentamicin resulted in little hair cell loss at 11h following washout, while loss was near completion at that time for 100 and 200 µM gentamicin, and at intermediate levels with 50 µM gentamicin treatment. Overall, there were significant differences between 25 µM, 50 µM and 200 µM treatments (Tukey’s multiple comparison, p<0.0001) but not between 100µM and 200µm treatments. These results demonstrate that the rate of delayed death is related to the concentration of gentamicin during the initial exposure.

**Figure 2.**
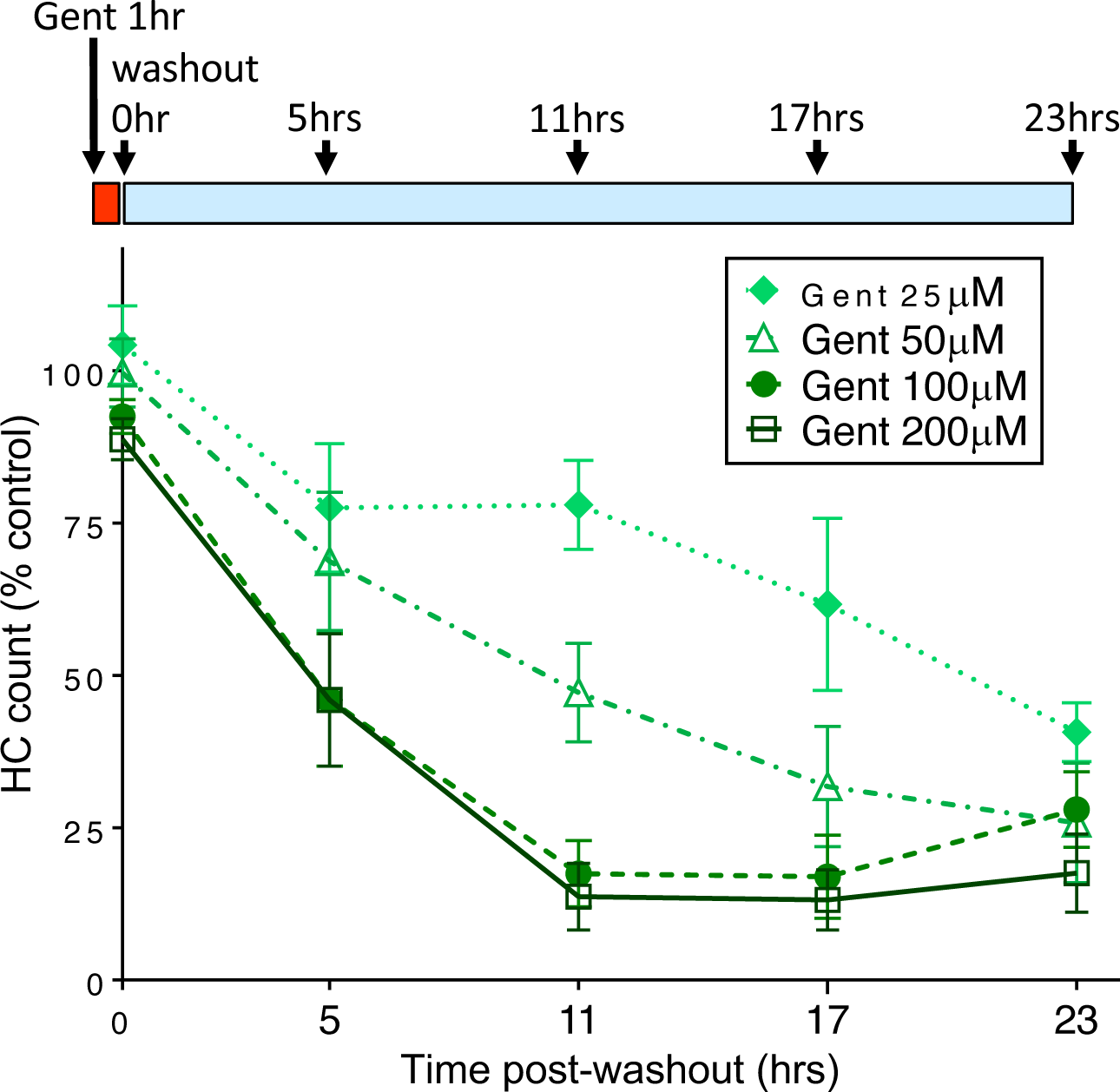
The rate of delayed hair cell loss is dependent on initial gentamicin concentration. Fish were treated with doses of gentamicin (25, 50, 100, 200 µM) for 1hr (red bar), then rinsed and incubated in fresh medium (blue bar). Loss of hair cells was assessed at 5hrs, 11hrs, 17hrs and 23hrs after the 1hr incubation period. Increasing initial dose results in more rapid delayed hair cell loss. Differences between 25 µM, 50 µM and 200 µM treatments were significant (Tukey’s multiple comparison, p<0.0001). There was no significant difference between 100 µM and 200 µm treatments. n=9-11 fish, 4 NMs/fish for each treatment. Error bars represent Standard Deviation.

Gentamicin is composed of a number of different structural AG subtypes that have differing levels of ototoxicity (O’Sullivan et al., 2020). We therefore tested whether we observed delayed death using G418, an AG closely related to gentamicin subtypes that can be readily obtained as a single, highly enriched isoform. As for gentamicin, we found that G418 had only a slight effect after 1h of exposure, while both 24h and 1+23 treatment paradigms resulted in substantial loss of hair cells (Supplemental Figure 1). This result suggests that differences in hair cell death over different concentrations and times of exposure are not caused by overlapping effects of distinct gentamicin isoforms.

### Different Ca^2+^ Dynamics During Acute and Delayed Hair Cell Death

We previously reported that acute neomycin exposure to zebrafish lateral line hair cells results in an increase in mitochondrial calcium followed by calcium increase in the cytoplasm, and that these events are necessary and sufficient to activate hair cell degeneration (Esterberg et al., 2013, 2014). To determine whether similar calcium changes occur during the delayed toxicity described above, we monitored intracellular calcium dynamics in hair cells using targeted, genetically-encoded calcium indicators, as performed previously (Esterberg et al., 2013; 2014). For these experiments we compared neomycin to G418 to avoid potential complexity of interpretation using the mixture of AGs in gentamicin. We used the green fluorescent GCaMP directed to mitochondria using a CoxVII targeting sequence (mitoGCaMP) and the red indicator RGECO localized to the cytoplasm (cytoRGECO) to measure compartmental calcium levels. We imaged cells under 3 different conditions: after exposure to 100 µM neomycin, where most cells undergo acute death, 100µM G418, where most cells undergo delayed death, or 400 µM G418, where some cells die under acute conditions and some die with a delayed time course. To capture acute response for treatment with 100 µM neomycin or 400 µM G418, we began imaging immediately after AG addition up to 1h. To capture delayed responses to treatment with 100 µM G418, fish were incubated in drug for 1h, rinsed and incubated a further 1.5h before imaging at 1 min intervals for 2h. For all treatments, hair cells may die at any time during the imaging period. Changes in fluorescence over time for individual cells were therefore aligned to the same endpoint of cell fragmentation.

Acute treatment with 100 µM neomycin resulted in increases in mitochondrial Ca^2+^ over baseline in all dying cells (16/16 dying cells; Figure 3A), consistent with previous results (Esterberg et al., 2014). By contrast, we rarely saw similarly large changes in mitochondrial Ca^2+^ when hair cells underwent delayed death after exposure to 100 µM doses of G418 (2/11 dying cells; Figure 3B). When treated with 400 µM G418, we observed numerous instances of cells undergoing increases in mitochondrial Ca2+ (10/14 dying cells; Figure 3C). The differences in proportion of cells showing a mitochondrial Ca^2+^ signal in each condition is highly significant (Chi-square = 20.25, p<0.0001). When we compared maximal changes in mitoGCaMP signal in dying cells under the three different AG exposure paradigms (Figure 3D), we found significant differences comparing treatment with 100 µM neomycin to 100 µM G418 (2.2±0.46 vs 1.2±0.32, Dunn’s multiple comparison test, p<0.0005). When we compared cytoplasmic Ca^2+^ changes using cytoRGECO (Supplemental Figure 2), we found a similar pattern: large changes in cytoplasmic calcium after acute 100 µM neomycin exposure (Supplemental Figure 2A), little or no changes after 100 µM G418 treatment (Supplemental Figure 2B), and intermediate changes in acute 400 µM G418 treatment (Supplemental Figure 2C). When we compared maximal changes in cytoRGECO signal in dying cells under the three different AG exposure paradigms (Supplemental Figure 2D), we found significant differences comparing treatment with 100 µM neomycin to 100 µM G418 (1.8±0.44 vs 1.2±0.38, Dunn’s multiple comparison test, p<0.0005). Taken together, these experiments further suggest that acute and delayed mechanisms of hair cell death have distinct underlying cellular mechanisms. Moreover, both types of mechanism can be induced by the same AG (here, G418) dependent on exposure concentration and time.

**Figure 3.**
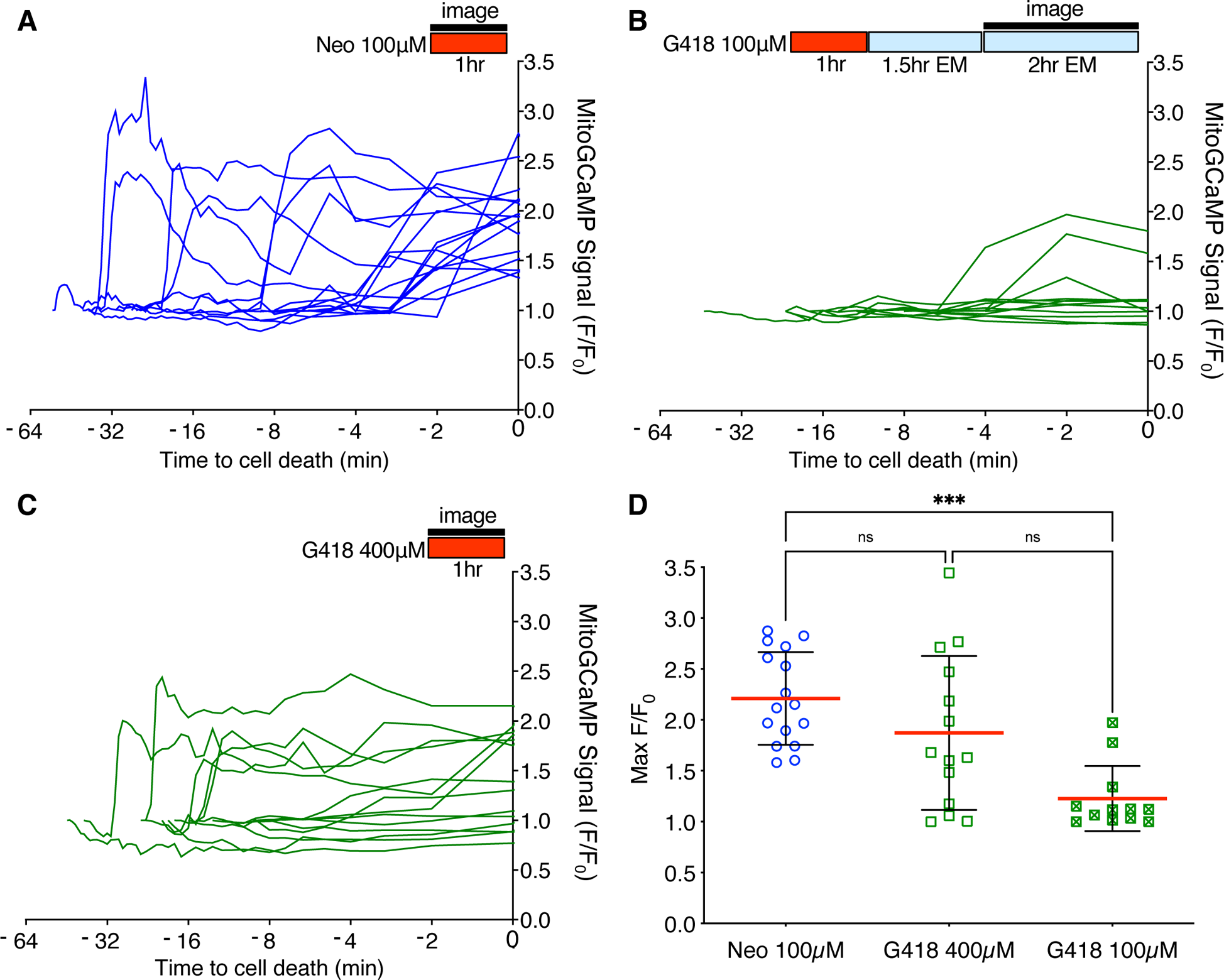
Different mitochondrial Ca^2+^ responses during acute or delayed hair cell death. A-C) Fish were incubated in AG (red bar) for time indicated, or rinsed into fresh medium (blue bar). Fluorescence changes above baseline (F/F_0_) from mitoGCaMP in response to AG addition were monitored at 30 sec intervals by spinning disk microscopy over the interval indicated (black bar). Individual traces represent responses of individual cells. Traces are aligned to time of cell fragmentation. A) Changes in mitoGCaMP signal in cells undergoing acute death in response to 100 µM neomycin. Hair cells were imaged during the first hour of neomycin exposure. Increases in mitochondrial Ca^2+^ were observed in 16/16 dying cells. B) Changes in mitoGCaMP signal in cells undergoing delayed death after exposure to 100 µM G418. Cells were exposed to G418 for 1hr followed by rinse and incubation in fresh embryo medium (EM) for 1.5hrs, and then imaged over an additional 2hr period. Increases in mitochondrial Ca^2+^ were observed in 2/11 dying cells. C) Changes in mitoGCaMP signal in cells undergoing acute death in response to 400 µM G418. Hair cells were imaged during the first hour of G418 exposure. Increases in mitochondrial Ca^2+^ were observed in 10/14 dying cells. D) Maximum mitoGCaMP signal compared to baseline for dying cells after neomycin or G418 exposure. ***Dunn’s multiple comparison test p<0.0005. Error bars represent Standard Deviation.

Mitochondrial calcium uptake after neomycin exposure results in production of reactive oxygen species (ROS), and treatment with mitochondria-targeted antioxidant mitoTEMPO partially protects hair cells from neomycin damage (Esterberg et al., 2016). We therefore tested whether mitoTEMPO could ameliorate effects of acute and delayed mechanisms of hair cell death (Figure 4). We found that 50 µM mitoTEMPO treatment conferred significant protection against 1h exposure to 100µM or 200µM neomycin (Figure 4A; Sidak multiple comparison test, p<0.0001), consistent with previous results. By contrast we observed no protection against 100µM or 200µM G418 using the 1+23h exposure paradigm (Figure 4B). These results support the idea that delayed cell death uses mechanisms that are distinct from those underlying acute cell death.

**Figure 4.**
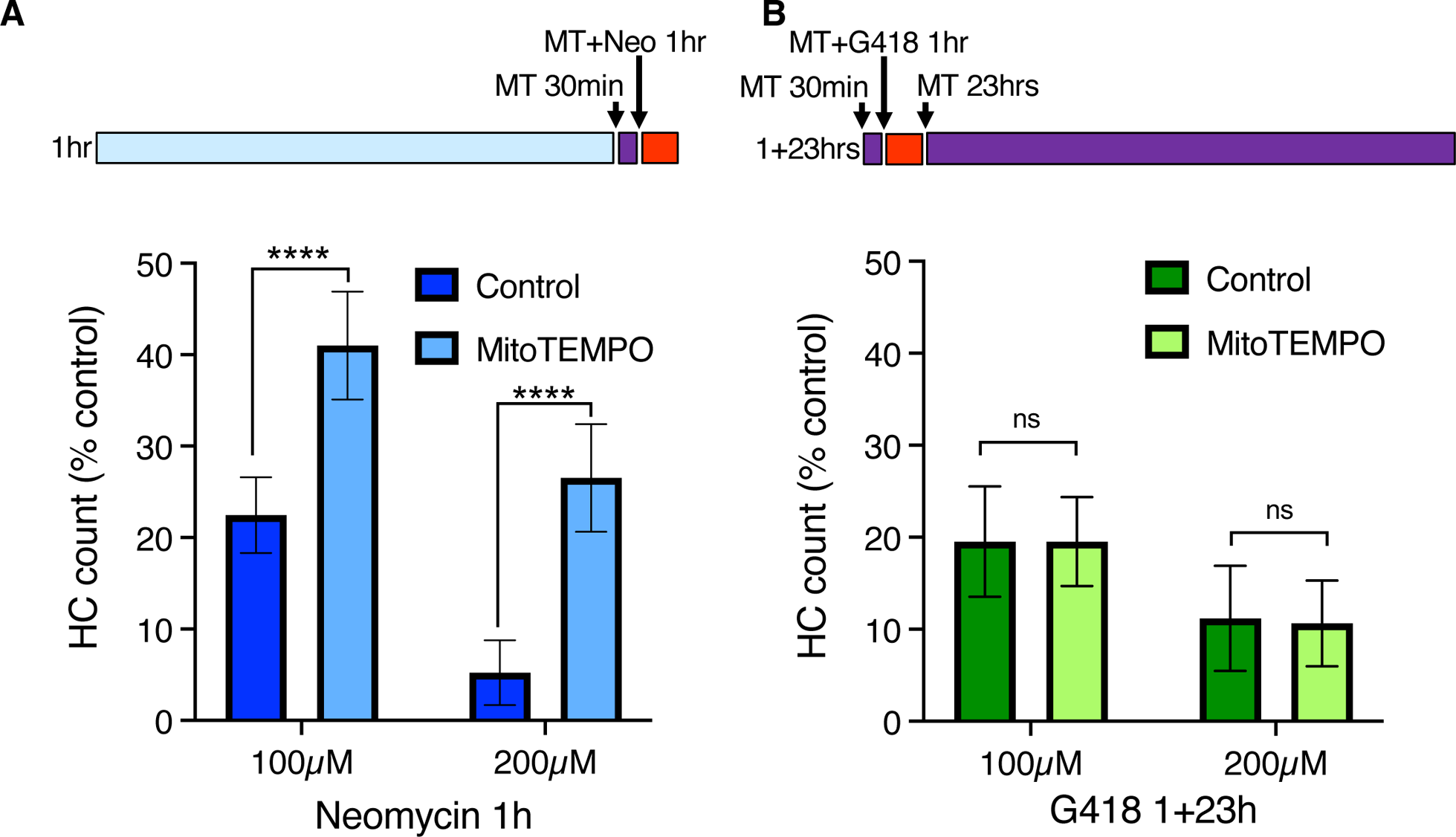
The mitochondrially-targeted antioxidant mitoTEMPO protects against acute neomycin damage from neomycin but not delayed damage from G418. 50 µM mitoTEMPO was added for 30min before AG (purple bar), co-treated with neomycin for 1hr or co-treated with G418 for 1hr (red bar), rinsed, and incubated with mitoTEMPO alone for 23hrs (purple bar). A) mitoTEMPO partially protected against damage from both 100 µM and 200 µM neomycin treatment. **** Two-way ANOVA, Sidak’s multiple comparison, p<0.0001. B) mitoTEMPO offered no protection against damage from either 100 µM or 200 µM G418 treatment. ns Two-way ANOVA, Sidak’s multiple comparison, p = 0.98. n=9-11 fish, 4 NMs/fish for each treatment group. Error bars represent Standard Deviation.

### Intracellular distribution of AG after washout

To begin identifying cellular pathways underlying the delayed death seen following gentamicin exposure, we examined more closely the cellular distribution of AGs at different times of exposure. For these experiments, we conjugated the red fluorescent dye Texas Red to neomycin (Neo-TR) or G418 (G418-TR) to follow entry and accumulation of AGs into hair cells. Neo-TR and G418-TR showed dose-dependent toxicity that was indistinguishable from the unconjugated forms (Supplemental Figure 2). To identify endolysosomal accumulation, we created a transgenic line where the small GTPase Rab7, a marker of late endosomes and lysosomes (Langemeyer et al., 2018), was conjugated to GFP and expressed under the hair cell-specific *myosin6b* promoter (Figure 5). We previously reported that neomycin showed broad cytoplasmic distribution while gentamicin was sequestered to puncta (Hailey et al., 2017). Here we see a similar pattern: after 5 minutes exposure, G418-TR is enriched within vesicles (Figure 5A), while Neo-TR shows diffuse distribution as well as some accumulation in Rab7-GFP+ vesicles (Figure 5B). To quantify the relative accumulation of AG in Rab7 vesicles, we used a semi-automated algorithm to segment the Rab7-GFP signal. We then measured Neo-TR or G418-TR fluorescence within the Rab7 compartment and compared it to TR fluorescence throughout the hair cell cytoplasm (Supplemental Figure 4). There was no significant difference between the number of vesicles segmented between the two conditions (Supplemental Figure 3B). Overall, the fluorescence signal was stronger in both compartments comparing Neo-TR to G418-TR (Supplemental Figure 3C-D), due to differences in labeling efficiency of each AG. When we compared the ratio of AG-TR fluorescence in Rab7 vesicles compared to the cytoplasm, we found that the ratio for G418-TR was significantly higher than that for Neo-TR (Figure 5C). These results suggest that differential accumulation of the distinct AGs into different compartments might underlie the distinct mechanisms of cell death observed after treatment.

**Figure 5.**
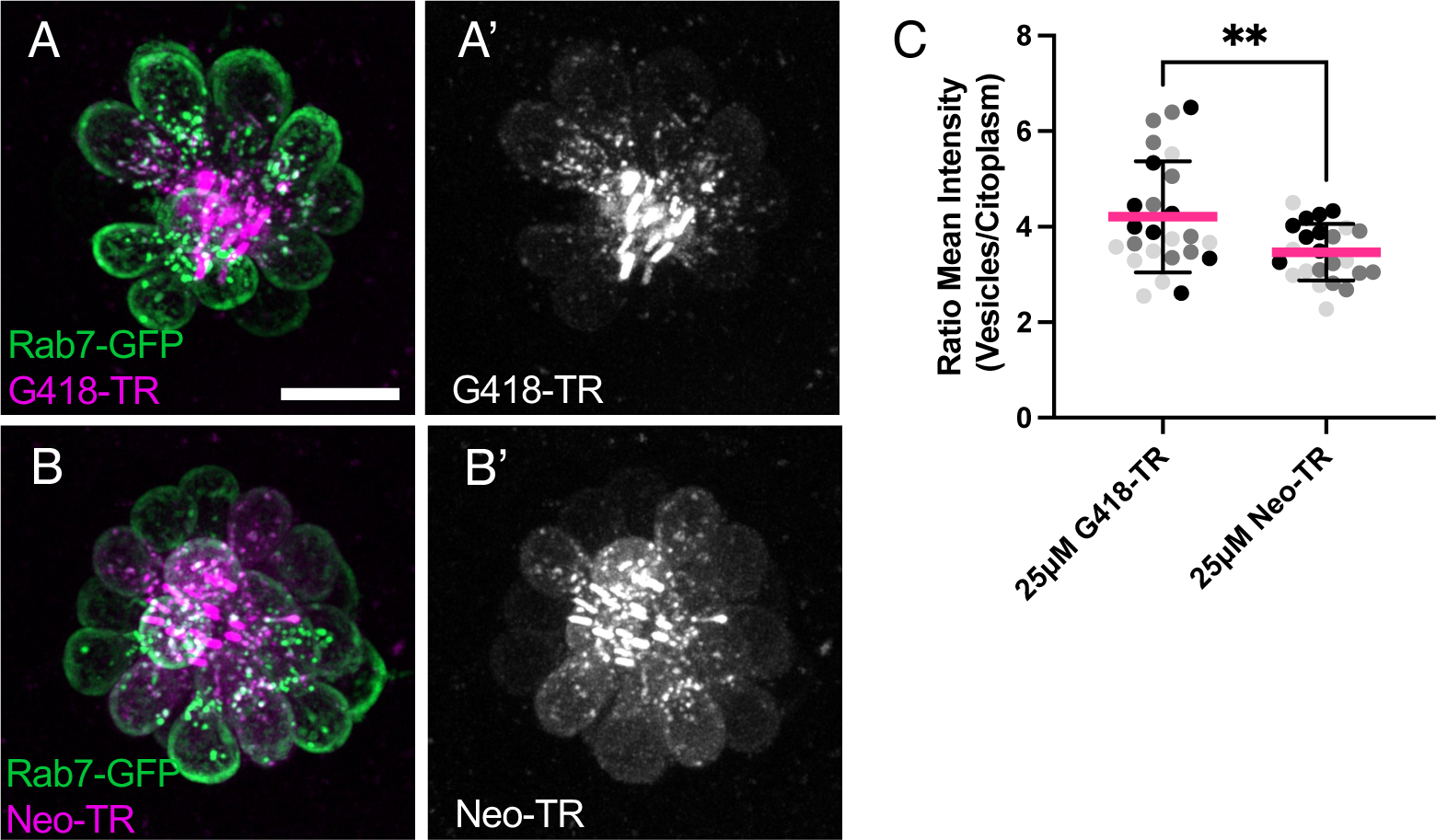
Neomycin and G418 differentially accumulate in Rab7+ vesicles. A) G418-TR (magenta) accumulation in Rab7+ vesicles (green) in neuromast from Tg(*myosin6b:EGFP-Rab7a)* transgenic line. A’) G418-TR signal is mainly in vesicles. B) Neo-TR (magenta) accumulation in Rab7+ vesicles (green) in neuromast from Tg(*myosin6b:EGFP-Rab7a)* transgenic line. B’) Neo-TR signal is found in vesicles and cytoplasm. C) Ratio of G418-TR and Neo-TR found in Rab7+ vesicles compared to cytoplasm. ** Unpaired T test, p<0.01. Error bars represent Standard Deviation.

### Altering endolysosomal function protects against delayed but not acute hair cell death

To further explore the idea that endolysosomal localization plays a central role in the different mechanisms of hair cell death, we treated hair cells with glycyl-L-phenylalanine 2-naphthylamide (GPN), a di-peptide that accumulates in lysosomes and disrupts their function (Jadot et al., 1984; 1990; Berg et al., 1994). To assess the effects of GPN on the distribution of G418, we measured the accumulation of G418-TR in vesicles in control animals or those pre-treated with 250 µM GPN for 1h and then co-treating with 50 µM G418-TR for 1h (Figure 6). We used semi-automated algorithms to segment hair cell bodies (Figure 6A’, B’) and G418+ puncta (Figure 6A’’, B’’). We found that GPN treatment significantly reduced the number of vesicles containing G418-TR (Figure 6C, Mann-Whitney test, p<0.0001). By contrast, the mean vesicle volume per neuromast (Figure 6D) or mean vesicle fluorescence (Figure 6E) remained unchanged. Moreover, the total fluorescence of G418-TR per neuromast was unchanged, suggesting that GPN affects the intracellular distribution of G418 but does not block uptake into hair cells.

**Figure 6.**
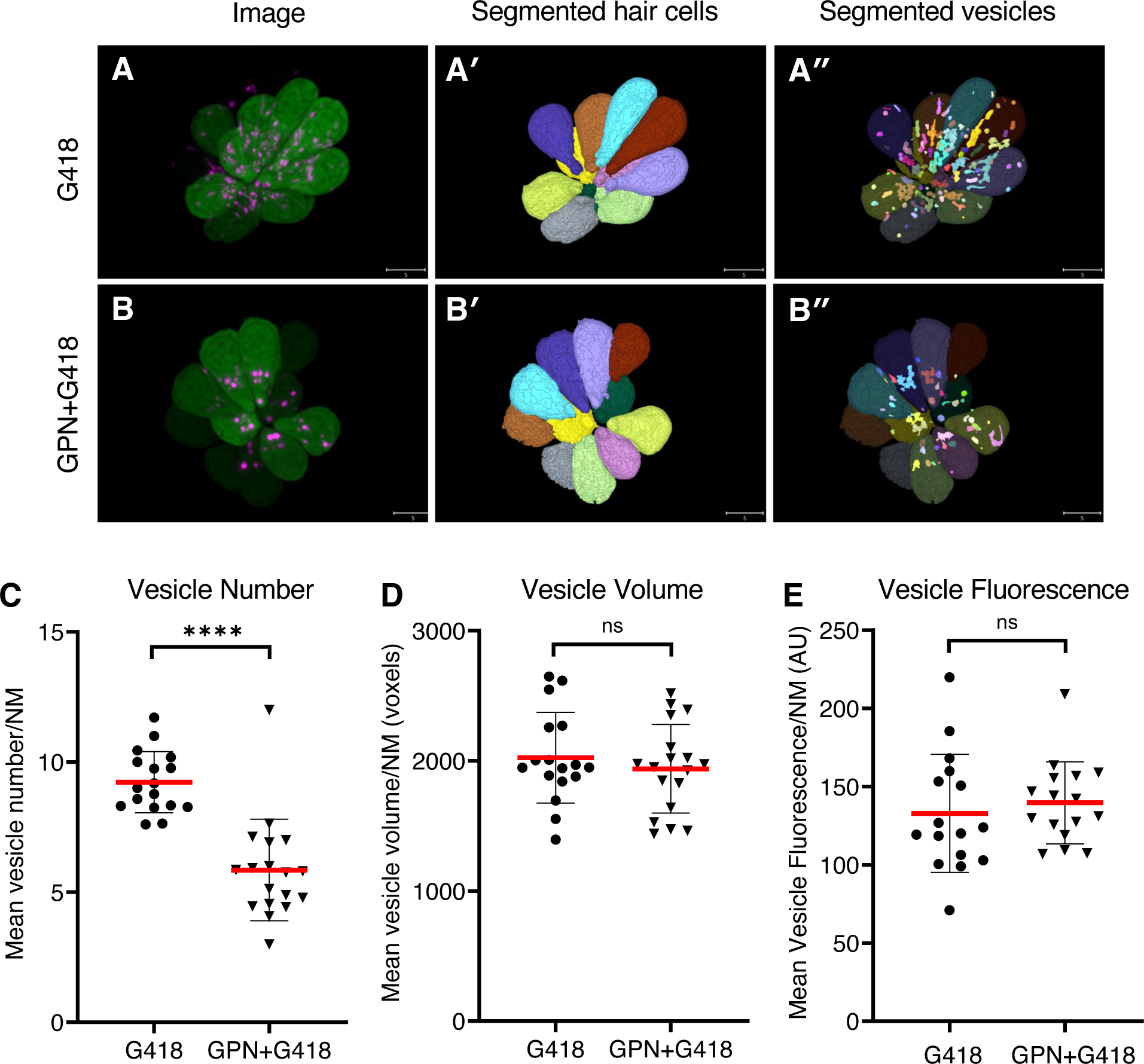
GPN treatment reduces the number of vesicles accumulating G418. A) Image of neuromast treated with G418-TR. A’) Mask of segmented hair cells. A’’) Mask of segmented vesicles. B) Image of neuromast treated with G418-TR and GPN. B’) Mask of segmented hair cells. B’’) Mask of segmented vesicles. C) Number of vesicles per hair cell, mean for each neuromast analyzed. ****Mann-Whitney test, p<0.0001. D) Vesicle volume per hair cell, mean for each neuromast. ns, Mann-Whitney test. E) Vesicle fluorescence per hair cell, mean for each neuromast analyzed. ns, Mann-Whitney test. Error bars represent Standard Deviation.

We next compared the effects of GPN treatment on toxicity due to exposure to neomycin or gentamicin. We assessed whether GPN altered delayed damage after gentamicin treatment by pre-treating with 250 µM GPN for 1h, co-treating with gentamicin at different concentrations, rinsing with fresh medium, and incubating in 250 µM GPN alone for an additional 23h (Figure 7A). GPN treatment offered robust protection from delayed death at all concentrations tested (Two-way ANOVA, Sidak multiple comparison, p<0.0001). We also assessed acute damage from neomycin after 1h GPN pre-treatment and 1h neomycin co-treatment (Figure 7B). Across all doses of neomycin, we found GPN treatment offered no protection. We also tested whether the vacuolar ATPase inhibitor bafilomycin A1, a well-characterized inhibitor of lysosomal function (Bowman et al., 1988), alters AG susceptibility (Supplemental Figure 5). Treatment with 100 nM bafilomycin A1 offered robust protection against gentamicin with significant differences at all concentrations (two-way ANOVA, Sidak multiple comparison, p<0.0001 for 25-100 µM, p<0.005 for 200 µM). Like GPN, bafilomycin offered no protection against neomycin. Finally, we tested whether treatment with these lysosomal drugs affected the overall uptake of fluorescent AG into hair cells (Supplemental Figure 6). Bafilomycin A1 significantly reduced G418-TR uptake (Mann Whitney test, p<0.0001). However, GPN treatment had no effect on the overall accumulation of G418-TR. Taken together, these results support a model where differential accumulation into the endolysosomal compartment leads to distinct mechanisms of hair cell death.

**Figure 7.**
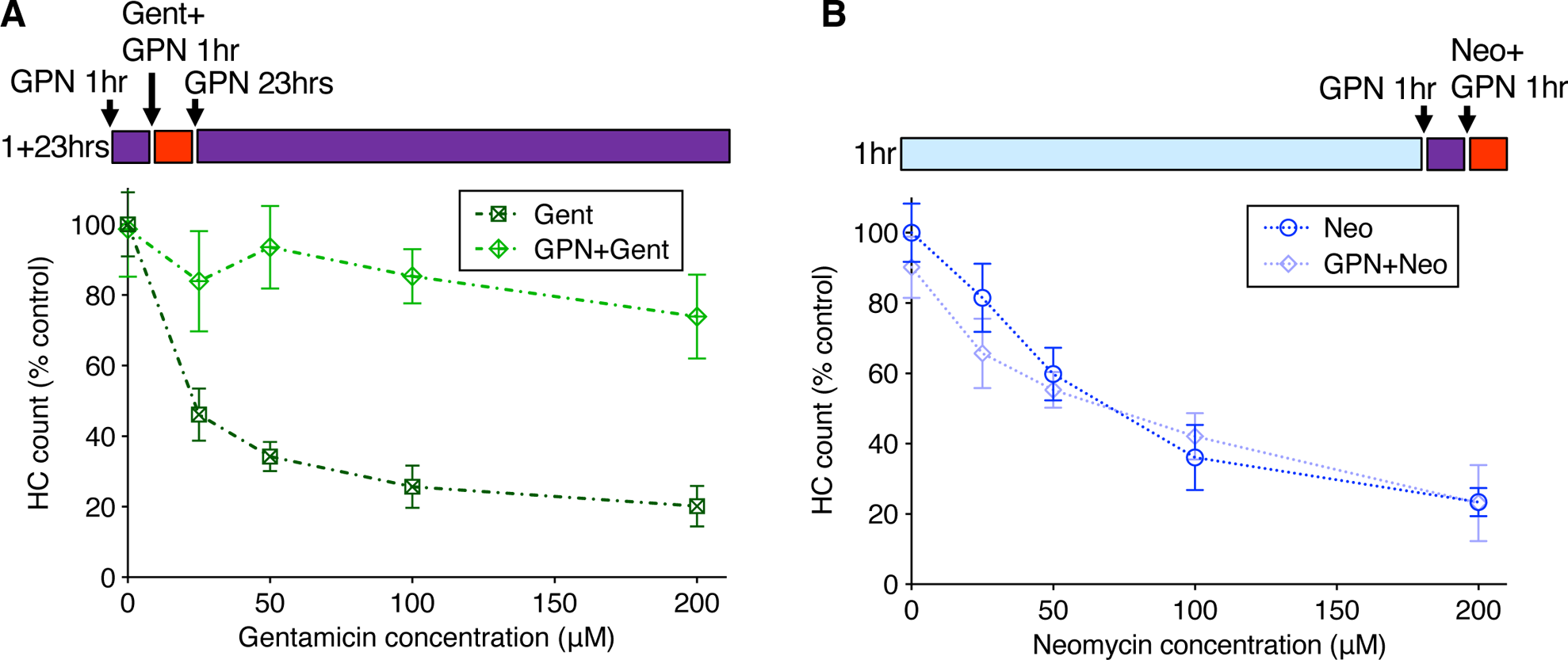
GPN treatment protects against delayed gentamicin damage but not acute neomycin damage. 250 µM GPN was added for 30min before AG (purple bar), co-treated with neomycin for 1hr or co-treated with G418 for 1hr (red bar), rinsed, and incubated with GPN alone for 23hrs (purple bar). A) GPN treatment offers robust protection against delayed death from all concentrations of gentamicin (Two-way ANOVA, Sidak’s multiple comparison, p<0.0001). B) GPN does not protect against acute death from neomycin at any concentration. n=9-11 fish, 4 NMs/fish for each treatment group. Error bars represent Standard Deviation.

## Discussion

In this study, we describe experiments that reveal at least two different mechanisms by which AGs kill zebrafish lateral line hair cells that can be distinguished by the time of exposure and the type of AG: we observed acute death within minutes of exposure to neomycin, but delayed death after washout exposure to gentamicin or G418. We provide evidence that these distinct mechanisms of acute or delayed death are regulated by different intracellular processes. We previously described an acute series of events through which a breakdown in inter-organellar Ca*^2+^* exchange results in mitochondrial collapse within minutes of neomycin exposure (Esterberg et al., 2013, 2014, 2016). Here we observed only rare instances of Ca*^2+^* increases in either cytoplasmic or mitochondrial compartments before cell fragmentation during delayed death after G418 exposure. Consistent with these results, the mitochondria-targeted antioxidant mitoTEMPO protected against acute death from neomycin but had little effect on delayed hair cell death from G418 exposure. We also demonstrated that using very high doses of G418 can result in acute death with accompanying Ca*^2+^* increases, indicating that the same AG can kill cells by acute or delayed mechanisms dependent on time and exposure concentrations. Finally, we show that endolysosomal accumulation of AG correlates with whether cells will undergo delayed death, and interfering with the endolysosomal compartment with the di-peptide drug GPN protects against delayed death but not acute death. Together, these results suggest that AGs engage different cell death pathways with distinct spatial and temporal dynamics.

The underlying mechanisms by which lysosomal accumulation of AGs result in delayed cell death remain unclear. We found that GPN treatment reduced the number of vesicles containing labeled G418 as well as protecting against delayed hair cell death, raising the possibility that sequestration is the cause of the delay, and subsequent release of AG from lysosomes results in hair cell death. However, differences in calcium dynamics accompanying acute and delayed death suggest that downstream events still diverge with the different AG treatments. Another possible mechanism is that lysosomal accumulation of AGs eventually promotes lysosomal membrane permeabilization (LMP) and cell death (reviewed in Wang et al., 2018). Several potential mechanisms lie downstream of LMP, including capthepsin-mediated cell death, changes in iron metabolism resulting in ferroptosis, or inflammasome activation resulting in pyroptosis. Work over the past decade has revealed that lysosomes function as more than just waste storage, but act as signaling hubs for nutrient sensing, lipid metabolism and intraorganellar contacts (Ballabio and Bonifacino, 2020); disruption of these processes by AGs may also contribute to hair cell death. GPN has long been used to disrupt lysosomal function (reviewed in Morgan et al., 2020), however the consequences of GPN treatment can vary depending on context. After its cleavage by lysosomal cathepsin proteases, GPN can promote LMP, altering lysosomal calcium signaling and pH (Yuan et al., 2021). GPN can also promote release of calcium from the ER, potentially independent of lysosomal pH changes (Atakpa et al., 2019). Future work will require genetic and pharmacological interventions to identify whether these pathways act downstream of lysosomal AG accumulation.

Our previous study suggested that AGs accumulate in lysosomes in zebrafish lateral line hair cells by autophagy, as treatment with the inhibitor 3-methyadenine (3MA) blocked vesicular localization (Hailey et al., 2017). 3MA was also effective at preventing AG-induced cell death in zebrafish (Coffin et al., 2009). In mouse cochlea, AG binding to the Rho-family interacting protein RIPOR2, activates autophagy and mitophagy via PINK/Parkin, and genetic inactivation of autophagy/mitophagy components protects hair cells from damage (Li et al., 2022). By contrast pharmacologic manipulation of autophagy suggests that it protects against AG induced hair cell death (He et al., 2017; Zhang et al., 2023). Screening a library of drugs that alter autophagy had mixed results, with some compounds offering protection while others showing toxicity (Draf et al., 2021). As a whole, these results suggest complex interactions between autophagy and AG toxicity, in part reflecting the underlying complexity of the autophagy network of signaling interactions.

Taken together, our work describes two distinct mechanisms of hair cell death in the zebrafish lateral line following two different routes through the cell, engaging either mitochondria or lysosomes. More broadly, many mechanisms have been previously described to underlie AG-induced hair cell death. These include caspase-dependent and caspase-independent apoptotic like processes (Forge and Li, 2000; Cunningham et al., 2002; Matsui et al., 2002, Jiang et al., 2006a), ribotoxic stress pathways (Francis et al, 2013) and necroptosis (Ruhl et al., 2019). AGs may damage hair cells by directly interfering with mitochondrial ribosomal translation (Prezant et al, 1993; Zhao et al., 2004; Hobbie et al., 2008; Matt et al., 2012; Shulman et al., 2014) or bioenergetics (O’Reilly et al., 2019b). AGs have also been described to preferentially interact with phosphoinositide lipids (Schacht, 1976), with multiple potential downstream consequences (Au et al., 1987; Priuska and Schacht, 1995; Lesniak et al., 2005, Jiang et al., 2006b). To what degree these processes intersect with the mitochondrial and lysosomal pathways remains an active area of interest.

We propose that engagement of different mechanisms of cell death depends on the type of AG, concentration, duration of exposure and potentially additional factors such as hair cell type, genetic factors and exposure to previous environmental insults. The potential for activation of multiple distinct cell death pathways to varying degrees in response to different AGs has important consequences for the design of therapeutic approaches to prevent AG-induced ototoxicity (reviewed in Kim et al., 2022 Hsieh, et al., 2024). It suggests that targeting specific downstream pathways may have limited success in preventing damage, as other mechanisms may bypass the intervention. This may explain the limited efficacy that blocking reactive oxygen species has had in preventing damage from ototoxins. It also suggests that approaches to develop compounds that block common early steps such as AG entry into hair cells (Kenyon et al., 2017; Chowdhury et al., 2018, O’Reilly et al., 2019a, de la Torre et al., 2024) or modifications of AGs to reduce hair cell uptake (Huth et al., 2015), may be promising directions for future therapy.

## Acknowledgments

We thank David White and the UW Zebrafish Facility for excellent fish care. This work was supported by a grant from the National Institutes on Deafness and Other Communication Disorders (R01 DC05987) to DWR.

