## Supplemental Information for "Multiple mechanisms of aminoglycoside ototoxicity are distinguished by subcellular localization of action"

**SIUPPLEMENTAL INFORMATION**

**Supplemental Figure 1.** G418 causes delayed hair cell death comparable to gentamicin. Dose-dependent loss of hair cells after treatment with G418 for 1hr, 24hrs or 1+23hrs. Differences between treatments were highly significant (2-way ANOVA, Tukey's multiple comparison, $p<0.0001$). n=9-11 fish, 4 NMs/fish for each condition.

**Supplemental Figure 2.** Different cytoplasmic calcium responses during acute or delayed hair cell death. Fluorescence changes above baseline ($F/F_0$) from cytoRGECO in response to AG addition, imaged by spinning disk microscopy at 30 sec intervals. Individual traces represent responses of individual cells. Traces are aligned to time of cell fragmentation. A) Changes in cytoRGECO signal in cells undergoing acute death in response to 100 μM neomycin. Hair cells were imaged during the first hour of neomycin exposure. Increases in mitochondrial $Ca^{2+}$ were observed in 10/10 dying cells. B) Changes in cytoRGECO signal in cells undergoing delayed death after exposure to 100 μM G418.  Cells were exposed to G418 for 1hr, followed by rinses and incubation in fresh embryo medium (EM) for 1.5h, and then imaged over an additional 2hr period. Increases in cytoplasmic $Ca^{2+}$ were observed in 2/16 dying cells. C) Changes in cytoRGECO signal in cells undergoing acute death in response to 400 μM G418. Hair cells were imaged during the first hour of G418 exposure. Increases in cytoplasmic $Ca^{2+}$ were observed in 4/12 dying cells. D) Maximum cytoRGECO signal compared to baseline for dying cells after neomycin or G418 exposure. ***Dunn's multiple comparison test $p<0.0005$. Error bars represent Standard Deviation.

**Supplemental Figure 3.** Texas Red label does not change efficacy of aminoglycosides. A) Comparison of G418 to G418-TR. There are no differences in dose-response relationships for either 1hr or 1+23hr treatments (Two-way ANOVA, Sidak's multiple comparison test). B) Comparison of neomycin to Neo-TR. There is no difference between dose-response relationships between unlabeled and labeled neomycin (Two-way ANOVA, Sidak's multiple comparison test).

**Supplemental Figure 4.** Automated segmentation of Rab7-labeled vesicles and neuromasts. A) Masks generated for vesicles, neuromasts, cytoplasm (neuromast-vesicle) and background. B) Ratio of vesicle area to neuromast area. There is no difference between drug treatment conditions (unpaired T test). C, D) Mean fluorescence values for whole neuromast (C) or vesicles (D). Error bars represent Standard Deviation.

**Supplemental Figure 5.** Bafilomycin A1 protects hair cells against G418 but not neomycin.
A) 100 nM Bafilomycin A1 treatment offers robust protection against gentamicin (Two-way ANOVA, Sidak's multiple comparison: ns, 10 μM; $p<0.0001$, 25, 50, 100 μM; $p<0.01$, 200 μM). B) Bafilomycin A1 does not protect against neomycin at any concentration. n=9-11 fish, 4 NMs/fish for each treatment group. Error bars represent Standard Deviation.

**Supplemental Figure 6.** Effects of GPN and Bafilomycin A1 on G418-TR uptake. A) 250 μM GPN treatment does not change G418 uptake. ns, Mann Whitney. B) 100 nM Bafilomycin A1 significantly reduces uptake. **** Mann Whitney, $p<0.0001$. n=12 fish, 6-7 NM/fish in each treatment group. Error bars represent Standard Deviation.

**Supplemental 1**

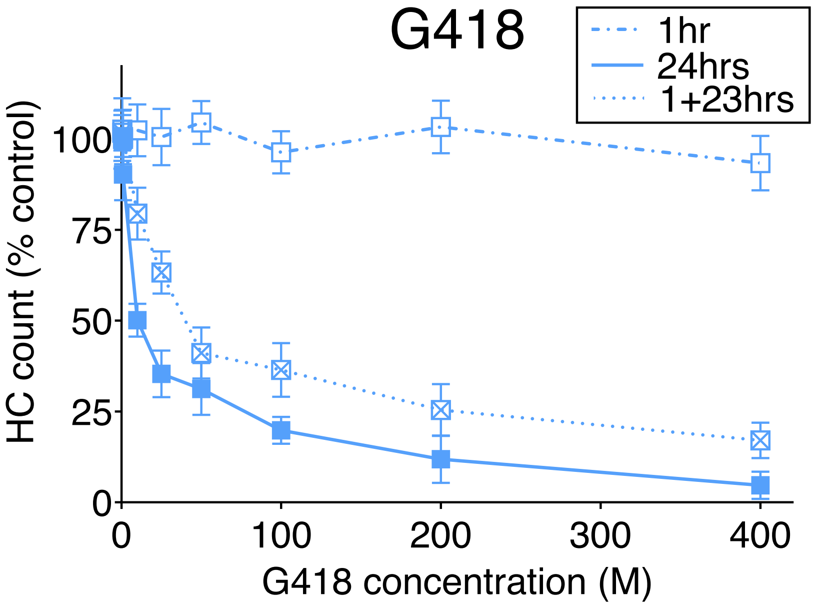

**Supplemental**
**Figure 2**

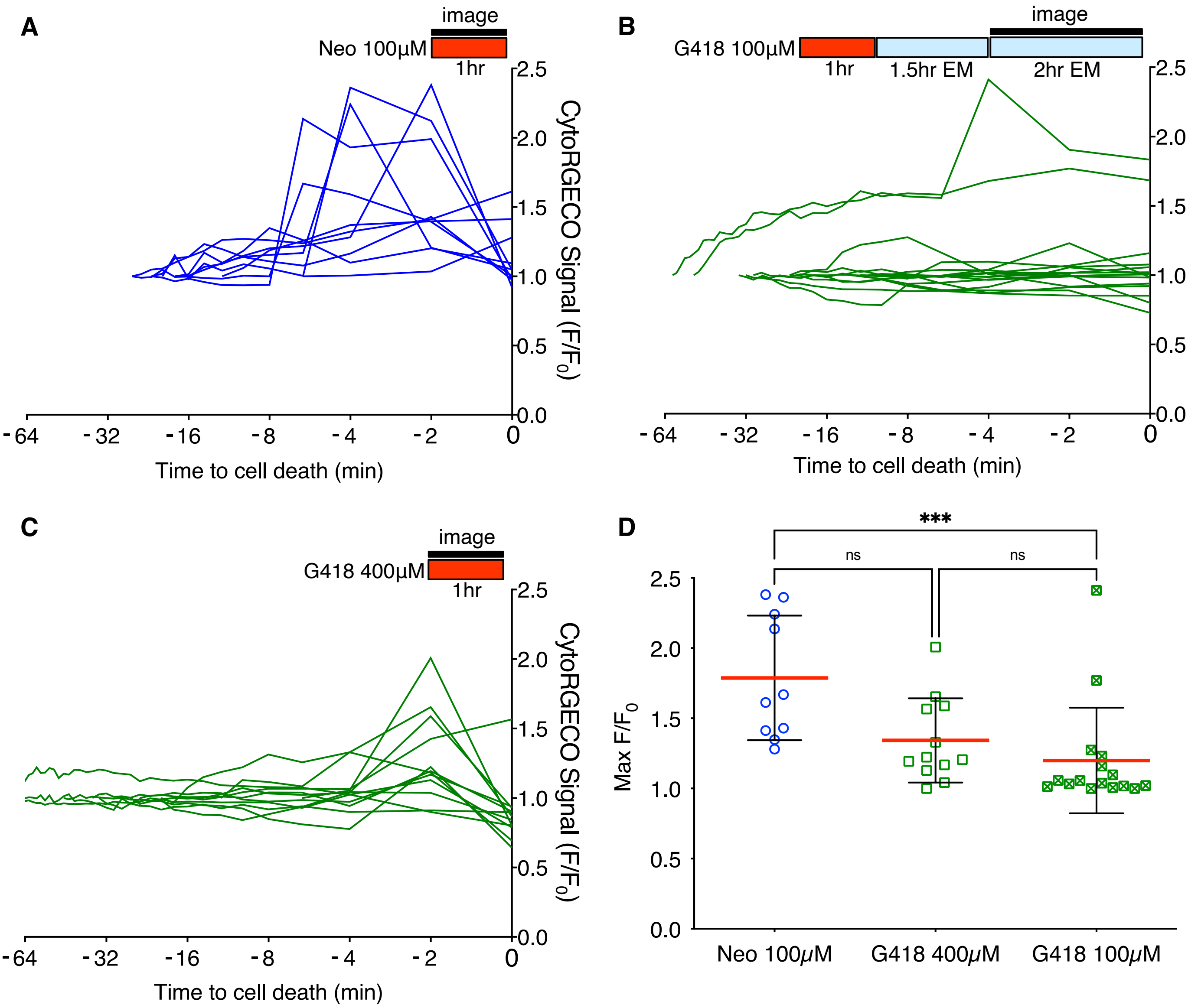

**Supplemental Figure 3**

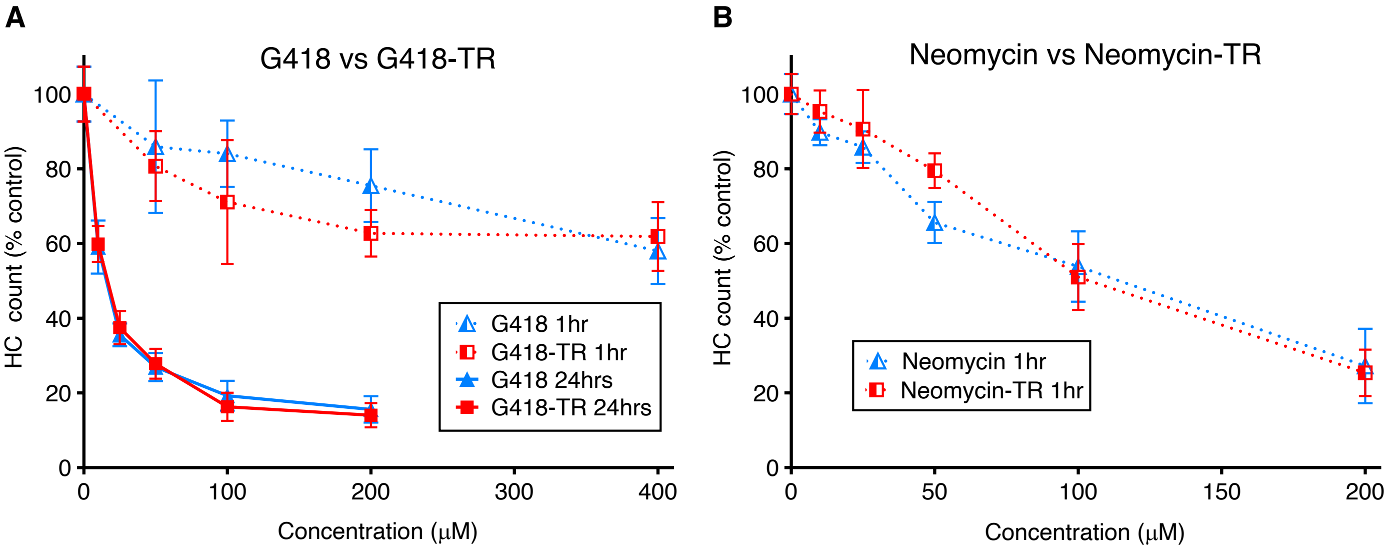

**Supplemental Figure 4**

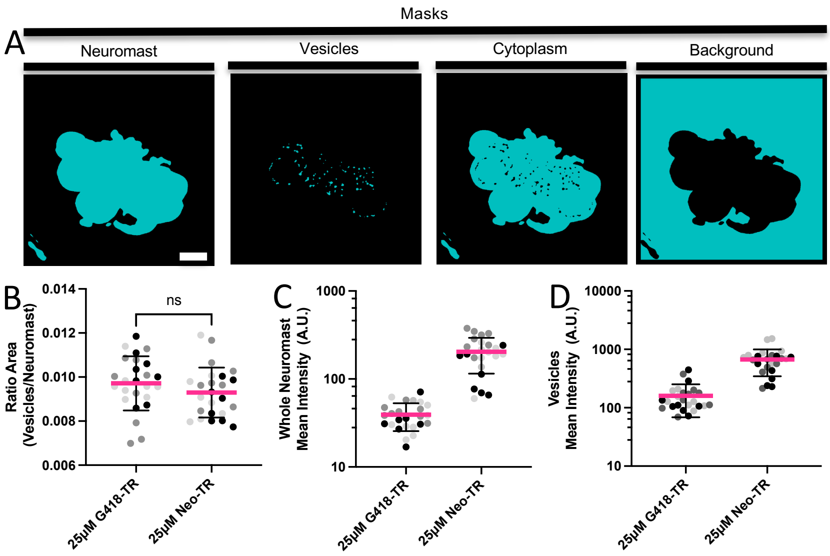

**Supplemental Figure 5**

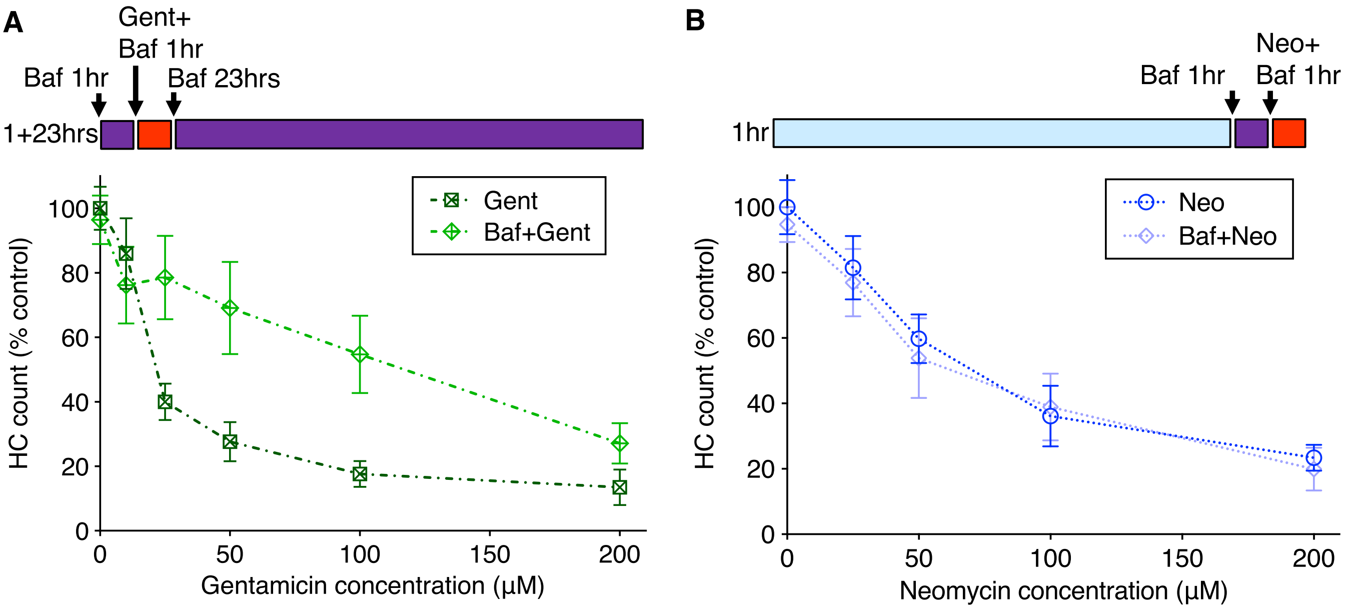

**Supplemental Figure 6**

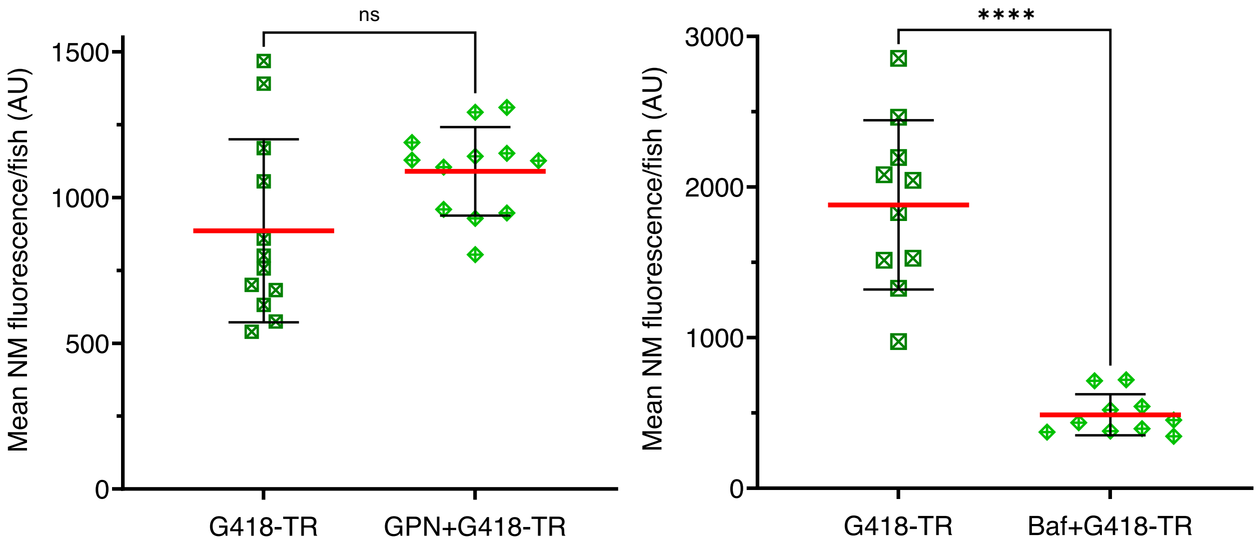
